# Switch-like activation of Bruton’s tyrosine kinase by membrane-mediated dimerization

**DOI:** 10.1101/284000

**Authors:** Jean K. Chung, Laura M. Nocka, Qi Wang, Theresa A. Kadlecek, Arthur Weiss, John Kuriyan, Jay T. Groves

## Abstract

The transformation of molecular binding events into cellular decisions is the basis of most biological signal transduction. A fundamental challenge faced by these systems is that protein-ligand chemical affinities alone generally result in poor sensitivity to ligand concentration, endangering the system to error. Here, we examine the lipid-binding pleckstrin homology and Tec homology (PH-TH) module of Bruton’s tyrosine kinase (Btk) Using fluorescence correlation spectroscopy (FCS) and membrane-binding kinetic measurements, we identify a self-contained phosphatidylinositol (3,4,5)-trisphosphate (PIP_3_) sensing mechanism that achieves switch-like sensitivity to PIP_3_ levels, surpassing the intrinsic affinity discrimination of PIP_3_:PH binding. This mechanism employs multiple PIP_3_ binding as well as dimerization of Btk on the membrane surface. Mutational studies in live cells confirm that this mechanism is critical for activation of Btk *in vivo*. These results demonstrate how a single protein module can institute a minimalist coincidence detection mechanism to achieve high-precision discrimination of ligand concentration.

## INTRODUCTION

In cellular decision-making, choices are often binary. However, the molecular input signals that serve as the basis for these decisions often rely on relatively small changes in ligand concentrations, occurring in a spatially and temporally fluctuating environment. One prominent example is the membrane-lipid second messenger phosphatidylinositol (3,4,5)-triphosphate (PIP_3_), which is involved in multiple signaling networks, including the PI3K/Akt/mTOR and T- and B-cell receptor pathways (Czech, 2000; Kane and Weiss, 2003). A wide variety of signaling proteins recognize PIP_3_ via pleckstrin homology (PH) domains. PH domain binding to PIP_3_ leads to membrane recruitment and activation of the signaling protein, typically by phosphorylation, and propagation of downstream signaling reactions (Lemmon, 2008; Lemmon and Ferguson, 2000; Newton, 2009). While this basic phenomenology is well described in many cases, it is also clear that simply relying on the ligand binding affinity would result in a gradual, hyperbolic response to the PIP_3_ density (Ferrell and Machleder, 1998; Papayannopoulos et al., 2005). Such a system would suffer a high probability of error as well as limited sensitivity, and would generally fall short of the signaling robustness exhibited in living cells.

One general mechanism to overcome this physical limitation, referred to as *coincidence detection*, relies on the binding of multiple membrane targets via distinct binding domains (Lemmon, 2008; McLaughlin et al., 2002; Newton, 2009). This mechanism is exemplified by the protein kinase C (PKC) family isozymes and the actin-nucleating WAVE complex (Hurley and Meyer, 2001; Johnson et al., 2000; Newton, 1995, 2009; Ziemba et al., 2014; Groves and Kuriyan, 2010). PKC has lipid binding modules that require independent binding to phosphatidylinositol-4,5-bisphosphate (PIP_2_), diacylglycerol (DAG), and Ca^2+^ for efficient membrane localization. Similarly, the WAVE complex undergoes a sequence of interactions with GTPases, acidic phospholipids, and PIP_3_ before activation (Chen et al., 2010; Kim et al., 2000; Lebensohn and Kirschner, 2009; Prehoda et al., 2000). In both cases, coincidence detection has the effect of producing spatial and temporal specificity, as well as a threshold-like response (Carlton and Cullen, 2005; Lemmon, 2008). Many of these examples have experimental foundations in structural data (Chen et al., 2010; Hurley and Meyer, 2001; Kim et al., 2000) and solution-based binding and activity assays (Johnson et al., 2000; Papayannopoulos et al., 2005; Prehoda et al., 2000). However, quantitative information, such as molecular stoichiometry, kinetic rates, and binding affinities, has largely been inaccessible for these membrane-bound processes. Thus, although some basic strategies for achieving nonlinear sensitivity and threshold-like discrimination are known, the precise molecular mechanisms and quantitative discriminating capabilities have not generally been experimentally established. Fortunately, there have been technological advances in quantitative biophysical methods, which, combined with membrane-mimetic models, have the potential to unmask these previously hidden processes (Chung et al., 2016; Groves and Kuriyan, 2010; Iversen et al., 2014; Lee et al., 2017; Lin et al., 2014; Ziemba et al., 2014).

In this work, the interaction between Bruton’s tyrosine kinase (Btk) and the membrane is analyzed by applying quantitative fluorescence spectroscopic and imaging methods to reconstituted systems. We focus on the PH-TH module of Btk, which consists of a classical PH domain fused to a zinc-bound Tec homology (TH) domain, a distinguishing feature of the Tec family kinases. Btk and other Tec family members resemble the Src and Abl families of non-receptor tyrosine kinases in that they contain Src homology 2 and 3 (SH2 and SH3) domains and a kinase domain (Figure 1B). The SH3-SH2 module plays a multifaceted role of suppressing the kinase before activation, localizing the protein to its target, and stabilizing the kinase after activation (Filippakopoulos et al., 2008; Nagar et al., 2003; Sicheri et al., 1997; Wang et al., 2015; Xu et al., 1997).

**Figure 1.**
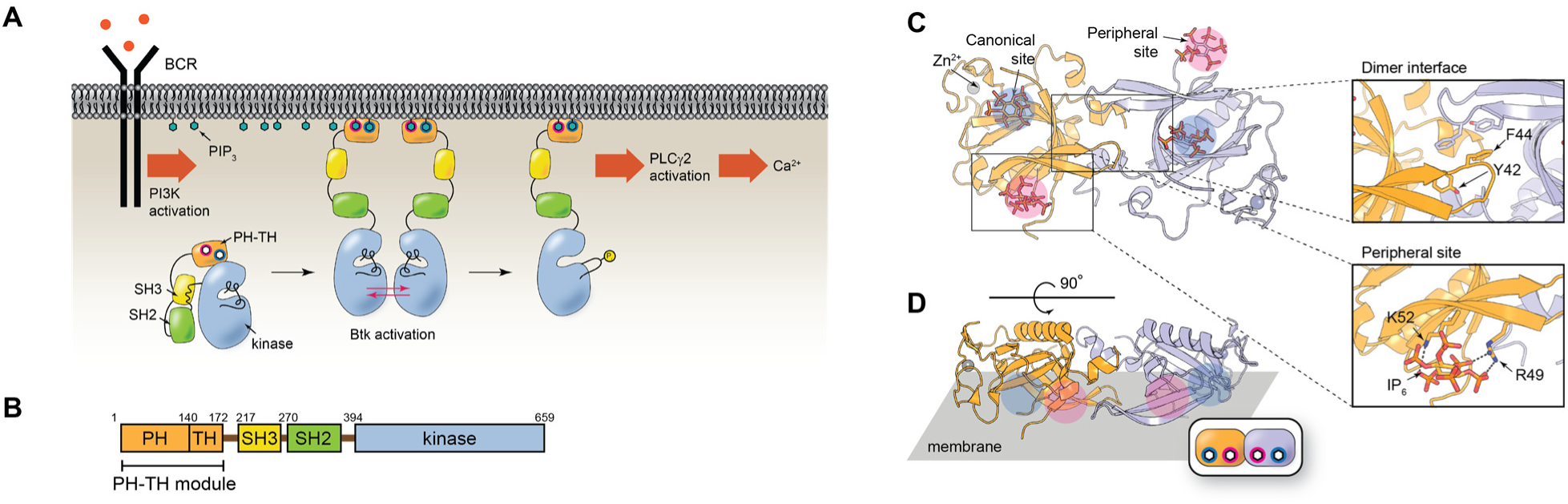
Btk in the B-cell receptor (BCR) pathway and the PH-TH domain structure. **(A)** Upon stimulation of BCR, PI3K produces PIP_3_, elevating the local PIP_3_ concentration. PIP_3_ recruits Btk to the plasma membrane, and Btk becomes activated it by *trans*-autophosphorylation, which then leads to PLCγ2 activation, calcium flux, and B-cell proliferation. **(B)** The domain architecture of Btk. **(C)** Schematic depicting the Saraste dimer of the Btk PH-TH domain with two IP_6_ coordinated to the canonical (blue) and peripheral (magenta) phosphoinositide binding sites (PDB 4Y94). Zoom-ins show the residues involved in the dimerization interface (top) and the peripheral inositol phosphate binding site (bottom) **(D)** Same structure rotated to show the orientation with respect to the membrane.

Btk plays a critical role in B-cell signaling, as depicted in figure 1A. When a B-cell encounters an antigen, the B-cell receptor promotes activation of phosphoinositide 3-kinase (PI3K). PI3K phosphorylates PIP_2_ to generate PIP_3_, which recruits Btk to the membrane via the PH-TH module. Btk is then activated, allowing it to activate phospholipase C-γ2 (PLC-γ2). PLC-γ2 hydrolyzes PIP_2_ into inositol (1,4,5)-trisphosphate (IP_3_) and diacylglycerol (DAG), leading to calcium flux and cellular activation (Hendriks et al., 2014; Scharenberg et al., 2007). Mutations in Btk have been shown to result in X-linked agammaglobulinemia, an autoimmune disease (Tsukada et al., 1993). Btk is the primary target of ibrutinib, which is used to treat some types of lymphoma and leukemia (Byrd et al., 2013; Hendriks et al., 2014).

Full-length Btk is maintained in an autoinhibited state in solution, but membrane recruitment releases the autoinhibition and leads to activation of Btk. Previous work with lipid vesicles has suggested that PIP_3_ strongly promotes Btk kinase activation via *trans*-autophosphorylation (Wang et al., 2015). While localization of a protein kinase to the membrane is expected to result in enhanced activity due to the local concentration effect (Grasberger et al., 1986), two key findings demonstrated that the PH-TH module may have a role in activation beyond simple membrane recruitment. First, Btk is strongly activated in solution by inositol hexaphosphate (IP_6_), a soluble inositol phosphate. In addition, a chimeric protein containing the PH-TH module of Btk artificially fused to the kinase domain of c-Abl is also activated by IP_6_. Second, crystal structures of the PH-TH domain (Figure 1C) reveal a dimeric structure (referred to as the Saraste dimer, as it was first identified by Matte Saraste and colleagues) (Baraldi et al., 1999b; Hyvonen and Saraste, 1997), and mutation of residues at this dimer interface prevents activation by IP_6_ in solution (Wang et al., 2015). These observations led to the hypothesis that there is a transient dimerization promoted by IP_6,_ and that it is important in Btk activation.

The crystal structure of the IP_6_:PH-TH complex revealed two IP_6_ binding sites on each PH-TH module, two of which form a Saraste dimer. One is the canonical binding site, common to other PH domains. The other is a secondary inositol phosphate binding site, referred to here as the peripheral site, that is proximal to the Saraste dimer interface (Figure 1B). Mutations at the peripheral binding site prevent the activation by IP_6_ of Btk, as well as the Btk-Abl chimera (Wang et al., 2015), suggesting that the occupation of the peripheral site by an inositol phosphate may allosterically influence the ability of the PH-TH module to dimerize (Baraldi et al., 1999a). However, dimerization of the Btk PH-TH module in solution has not been detected directly, with or without IP_6_. Also left open is the question of whether IP_6_ is a physiologically relevant ligand for the Btk PH-TH module, or whether it is mimicking the action of another ligand, such as PIP_3_, presented in the context of the plasma membrane.

In this paper, we demonstrate that PIP_3_ triggers the dimerization of the Btk PH-TH module on membranes in a switch-like manner that is dependent on the Saraste dimer interface. Critical for this behavior is the ability of the Btk PH-TH module to bind to two PIP_3_ lipids, creating a nonlinear response that is fourth-order with respect to the PIP_3_ surface density. This represents a minimalist coincidence detection mechanism, achieved with a single protein module. Further, we establish the physiological importance of the peripheral binding site and the Saraste dimer interface in the activation of the B-cells through activity assays performed in live B-cells. These results illustrate how the PH-TH domain of Btk takes advantage of multiple interactions to attain switch-like decision-making in B-cell signaling, and the simplicity of this mechanism suggests that it may be widespread in cellular signaling networks.

## RESULTS AND DISCUSSIONS

### The Btk PH-TH module dimerizes on membranes

Every crystal structure of the Btk PH-TH module shows the Saraste dimer, even when there is no ligand bound (Hyvonen and Saraste, 1997; Wang et al., 2015). Nevertheless, dimerization has not been detected directly in solution, suggesting that it is very weak. The membrane might, however, play a role in dimerization of the Btk PH-TH module, by increasing the local concentration or enforcing a specific orientation with respect to the lipid headgroups. To address this, we monitored the density-dependent diffusion of Btk on supported lipid bilayers (SLBs) containing PIP_3_. The experimental setup is shown in Figure 2A. eGFP-tagged Btk PH-TH module (henceforth referred to simply as PH-TH module) is introduced to glass-supported lipid bilayers containing PIP_3_ and a trace amount of Texas Red (TR) dye-labeled lipids. The dual-color excitation allowed the simultaneous measurement of the proteins and lipids by fluorescence correlation spectroscopy (FCS) or time-correlated single photon counting (TCSPC). TR lipid serves as a reference for each measurement, and the protein diffusion coefficient values reported for the PH-TH module here are shown relative to that of TR lipids measured simultaneously in the same spot.

**Figure 2.**
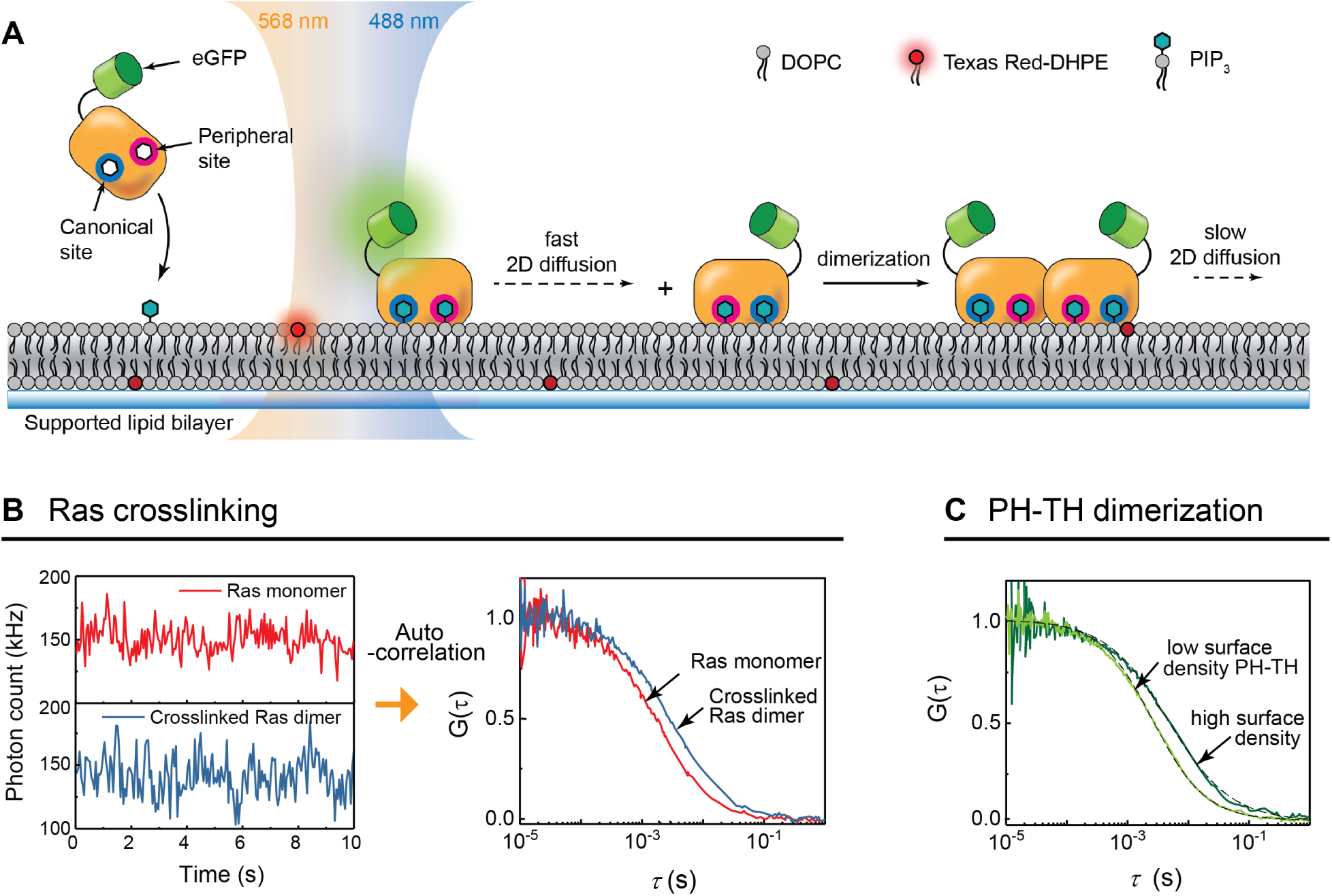
Detection of two-dimensional dimerization reaction on membrane surfaces by FCS. **(A)** In a dual-color FCS setup, Texas Red-labeled lipid (TR-DHPE) and eGFP-labeled PH-TH domain adsorbed to the SLB by PIP_3_ are simultaneously measured. **(B)** For membrane-bound Ras, time-dependent fluorescence intensity fluctuation due to diffusion is recorded (left), before (red) and after the addition of the RBD-LeuZ crosslinker (blue). The corresponding autocorrelation functions are shown on the right. **(C)** In the case of the PH-TH domain dimerization reaction, the difference in diffusion can be clearly resolved as the surface density is increased, the dimer population increases, which manifests as slower average diffusion rate.

In FCS, the time-dependent fluorescence intensity fluctuations due to fluorescent particles entering and exiting the focus area is recorded, as shown in Figure 2B (left). The autocorrelation function of the intensity fluctuations (right) has a decay profile with a correlation time constant, *τ*_d_, which is the average particle residence time in the focus area (Lakowicz, 2006). In this setup, the dominant process determining *τ*_d_ in the millisecond timescale is the lateral diffusion of the fluorescent species. FCS has been shown to be useful in measuring membrane-bound protein diffusion, and, in particular, for the detection of low-affinity dimerization on membranes (Chung, 2018).

Supported membranes are highly homogeneous and proteins bound to them exhibit unencumbered Brownian motion, which enables lateral diffusion to be used as a robust indicator of dimerization. The diffusion rate of a membrane-bound protein is determined by multiple factors, such as the fluidity of the surrounding environment, the shape of the protein, the geometry of the lipid anchors (Saffman and Delbruck, 1975), and any associations between proteins. In general, however, a peripheral membrane protein such as the Btk PH-TH module displays a mobility that is comparable to the lipids in which it is entrained, as the friction introduced by lipid anchors plays the dominant role in determining the overall diffusion of the protein. An association between membrane-bound proteins with each other generally results in a substantially slower diffusion (Chung, 2018; Knight et al., 2010). To illustrate the diffusion change on dimerization, the autocorrelation functions of Ras, a lipid-anchored protein, before and after crosslinking are shown: the residence time *τ*_d_ changes from 1.9 to 2.6 ms upon near-complete crosslinking, which corresponds to the reduction of diffusion coefficient from 4.2 to 2.3 μm^2^/s (data for Ras are taken from (Chung, 2018)). A similar change, from 3.5 to 1.7 μm^2^/s, is observed for the diffusion constant of the wild type Btk PH-TH module when the Btk surface density is increased, suggesting dimerization. Figure 2C shows the normalized autocorrelation functions for a lower surface density (20 μm^−2^, light green) and a higher surface density (400 μm^−2^, dark green) of the PH-TH module on the membrane.

The formation of Btk dimers on the membrane surface is a two-dimensional binding reaction whose binding free energy is reflected in a two-dimensional dissociation constant:

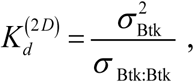

where *σ*_Btk_ and σ_Btk:Btk_ represent the two-dimensional surface density of Btk monomer and dimer, respectively. Thus, as in a three-dimensional binding titration, the equilibrium fraction of Btk dimers is a function of Btk monomer surface density. As a general strategy in these experiments, we monitor dimerization as a function of surface density to obtain the two-dimensional dimerization binding curve. A powerful aspect of the FCS measurement is that, in addition to mobility, it also provides a readout of surface density in the value of the autocorrelation function, *G*(0) = 1/*N*, where *N* is the number of particles at the focus area. Throughout this paper, Btk mobility and surface density data are determined from FCS measurements for each data point.

The surface density-dependent diffusion demonstrates that the wild type Btk PH-TH module is capable of dimerization on the membrane, shown in the left panel of Figure 3A (black solid circles). The mobility of monomeric PH-TH module bound to membrane via PIP_3_ is approximately 0.85 relative to TR-labeled lipid, as indicated by the diffusion coefficient at low surface densities. On a 4% PIP_3_ bilayer, the diffusion of the wild type PH-TH module at high surface densities approaches 0.4 relative to TR-labeled lipid diffusion, indicating an appreciable population of slowly diffusing dimers. The dimerization of the PH-TH module was further corroborated by density-dependent FRET, in which eGFP and mCherry labels on the Btk PH-TH modules were used as the donor and acceptor. The FRET efficiency on 4% PIP_3_ SLBs increases with the surface density, which is also consistent with dimerization (Figure 3C). Notably, these SLB-based measurements are the first direct evidence for Btk dimerization on membranes.

**Figure 3.**
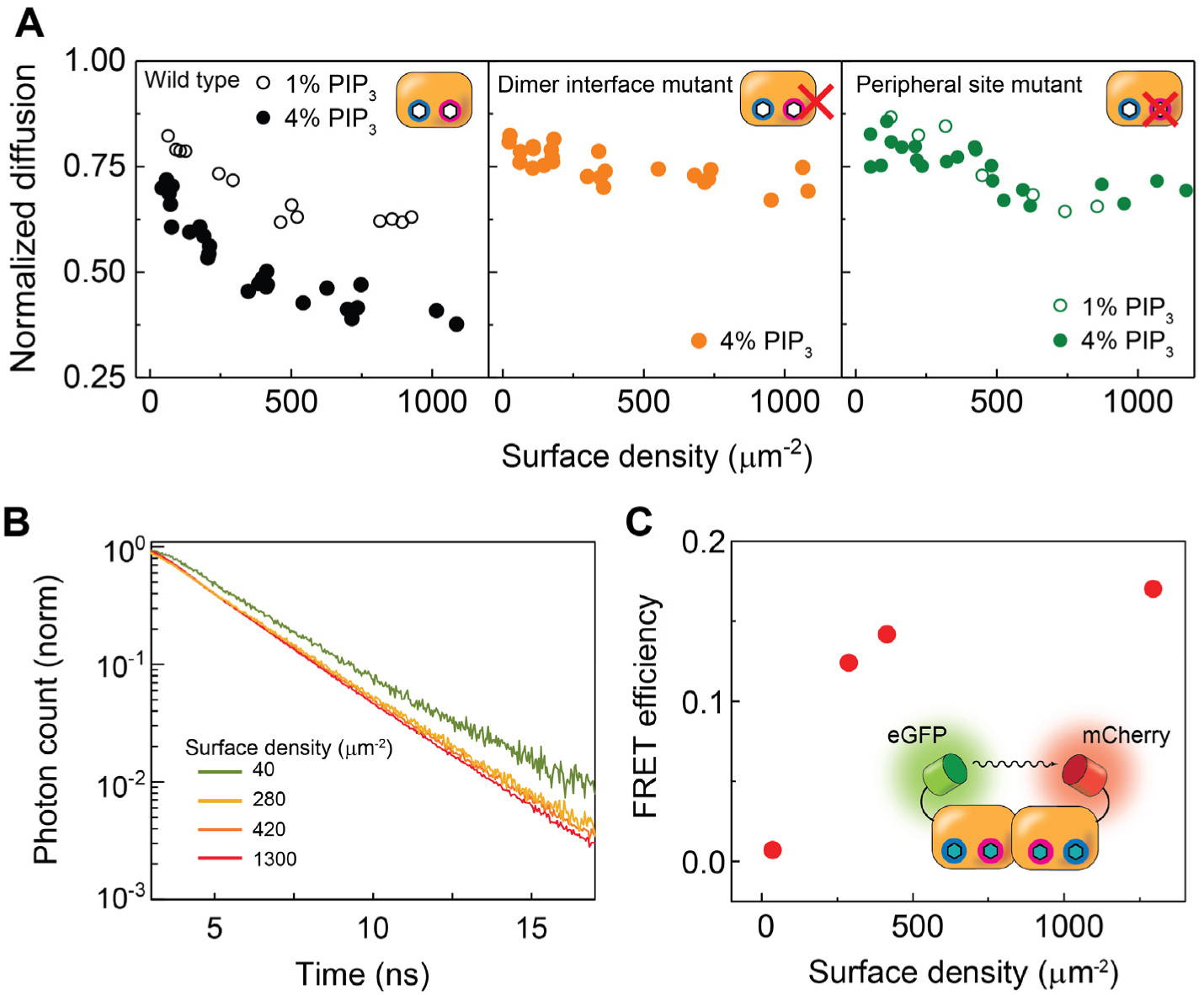
Diffusion and FRET measurement. **(A)** The density-dependent diffusion was measured for the wild type (left), dimer interface mutant (middle), and the peripheral site mutant (right) on SLBs containing 1% (empty circles) and 4% PIP_3_. In this set, only the wild type on 4% PIP_3_ bilayers shows a significant decrease in diffusion, which indicates dimerization. **(B)** The fluorescence lifetime of eGFP tagged to wild-type PH-TH module in presence of mCherry-labeled wild-type PH-TH modules was measured as a function of surface density on 4% PIP_3_ SLBs. **(C)** FRET efficiency increases as a function of the surface density, consistent with dimerization.

### Multiple PIP_3_ binding makes the PH-TH module sensitive to the PIP_3_ surface density

To examine how the membrane density of PIP_3_ affects Btk PH-TH dimerization behavior, protein diffusion was measured on supported lipid bilayers containing 1% and 4% PIP_3_. Previous measurements of Btk activity on PIP_3_ containing vesicles had shown an ultrasensitive activation occurring between 2 and 5% PIP_3_ (Wang et al., 2015). Therefore, we focused on this range, to see whether a change in dimerization could be detected. The left plot in Figure 3A shows diffusion measurements for 1% and 4% PIP_3_ bilayers. Compared to 4% PIP_3_ bilayers (filled black circles), the PH-TH module displays a significantly diminished potential for dimerization on 1% PIP_3_ bilayers (empty black circles) at comparable protein surface densities.

The fact that the PIP_3_ surface density modulates the dimerization capability of the PH-TH module at similar protein surface densities on the membrane leads to two points. First, each PH-TH module interacts with multiple PIP_3_ lipids: if there were a 1:1 stoichiometric relationship between the PH-TH module and PIP_3_, then its behavior would be identical regardless of the PIP_3_ density, as long as the protein surface density is constant. However, this is clearly not the case, meaning that there are additional PIP_3_ binding. Second, the additional PIP_3_ may be allosterically involved in a structural change or electrostatic interaction necessary for dimerization. Molecular dynamics simulations suggest that the PH-TH module can switch between conformations with different exposures of the dimer interface, and this equilibrium may be shifted by inositol binding at the peripheral site (Wang et al., 2015).

A remarkable feature in the PIP_3_ density-dependent dimerization is an ultrasensitive response to the PIP_3_ concentration. For example, according to estimates based on the FCS data, the surface density of the PH-TH module required for 15% dimer fraction is 1,110 μm^−2^ on 1% PIP_3_ and 170 μm^−2^ on 4% PIP_3_ (see SI 2 for details). The sharpness of the transition in dimerization behavior can be appreciated by considering that the solution concentrations necessary to achieve these membrane surface densities of the PH-TH module are 34 nM and 0.9 nM for 1% and 4% PIP_3_ bilayers, respectively, a 38-fold difference.

We also examined the specificity of the Btk PH-TH module towards PIP_3_ for recruitment and dimerization. Other PH domains are known to interact with negatively charged lipids with low selectivity (Kavran et al., 1998; Lemmon, 2008). Previous solution-based studies have shown that the Btk PH domain is highly specific for PIP_3_ over other types of anionic lipids (Rameh et al., 1997; Wang et al., 2015). Accordingly, no recruitment was observed in SLBs containing 10% PS or 4% PIP_2_. When a mutation was introduced such that a residue near the canonical binding site of the PH domain, Glu 41, which confers selectivity for PIP_3_ over PIP_2_ (Pilling et al., 2011), is removed, then there is significant recruitment of the mutant PH-TH module to PIP_2_-containing membranes (Figure S2). These observations suggest that the initial binding of the PH-TH module to the membrane is through specific PIP_3_ binding to the canonical lipid binding site rather than due to nonspecific charge interactions. Furthermore, inclusion of negatively charged PS does not promote dimerization in 1% PIP_3_ bilayers, indicating that dimerization occurs through specific interactions with PIP_3_.

### Mutations identify critical sites for dimerization

For this study, two double mutants were created based on the crystal structure of the PH-TH module bound to soluble IP_6_: a Saraste-dimer interface mutant (Y42Q/F44Q), and a peripheral site mutant (K49S/R52S). The mutated residues are depicted in the crystal structure of PH-TH module coordinated to two IP_6_ molecules (Figure 1C). The Saraste dimer interface mutation, highlighted in orange, disrupts dimerization by removing tyrosine and phenylalanine residues that provide intermolecular hydrophobic contacts (Baraldi et al., 1999a). The peripheral site mutation, in green, interferes with peripheral site binding by replacing lysine and arginine residues by serine (Wang et al., 2015). These mutations have been shown previously to impede Btk activation by IP_6_ in solution-based biochemical assays. However, a direct connection between peripheral site binding of lipids, dimerization, and phosphorylation had not been clarified. Here we sought to establish whether PIP_3_ can promote formation of the Saraste dimer by engaging the peripheral site of the PH-TH module.

The middle plot of Figure 3A shows the density-dependent diffusion of the dimer interface mutant. The diffusion remains relatively fast up to a high surface density, indicating that this mutation impairs dimerization. In light of the biochemical activity measurements in which this mutant is incapable of activation, the natural interpretation is that dimerization is required for activation via *trans*-autophosphorylation. The FCS diffusion measurements of the dimer interface mutant clearly establish the relevance of the crystallographically observed Saraste dimer for the membrane-bound dimerization we observe.

PIP_3_ binding at the peripheral site was also found to be necessary for dimerization, as mutation of this site resulted in a weaker density-dependent diffusion change on a 4% PIP_3_ bilayer (solid green circles, Figure 3A, right) compared to the wild type. Furthermore, the diffusion behavior is nearly identical on 1% PIP_3_ bilayers (empty green circles)—in other words, PIP_3_ concentrations cannot be differentiated without the peripheral site. Unlike the canonical phosphatidylinositol binding site, which is common to many PH domains, the role of the peripheral site in PIP_3_ binding had been ambiguous. There had been no direct evidence for PIP_3_ binding at this site; the soluble form of the PIP_3_ headgroup, inositol (1,3,4,5)-tetraphosphate (IP_4_), does not bind there when crystallized (Wang et al., 2015). Our new observations suggest that engagement of the peripheral site is important for the dimerization and activation of Btk on the membrane.

### Adsorption kinetics reveals dimerization driven by multiple PIP_3_ binding

The diffusion measurements of PH-TH module on membranes indicated that the Btk PH-TH module undergoes at least two PIP_3_ binding and dimerization, suggesting that the adsorption kinetics would be complex due to these multiple processes. The time-dependent adsorption profiles for wild type and mutant PH-TH constructs onto 4% PIP_3_ bilayer, monitored by total internal reflection fluorescence (TRIF) microscopy, are shown in Figure 4A. It is immediately apparent that compromising either the dimerization interface or the peripheral binding site has a significant impact on the protein-membrane interactions. The comparison of adsorption profile normalized to the final density shows that the dimer interface mutant is faster to reach equilibrium compared to the wild type, and that the peripheral site mutant is even faster (Figure 4B). This is consistent with the notion that the wild type PH-TH module undergoes more kinetic processes (PIP_3_ binding at the canonical and peripheral sites, and dimerization) than the dimer interface mutant (which can undergo PIP_3_ binding at the canonical and peripheral sites), or the peripheral site mutant (which binds lipid at the canonical site only and fails to dimerize).

**Figure 4.**
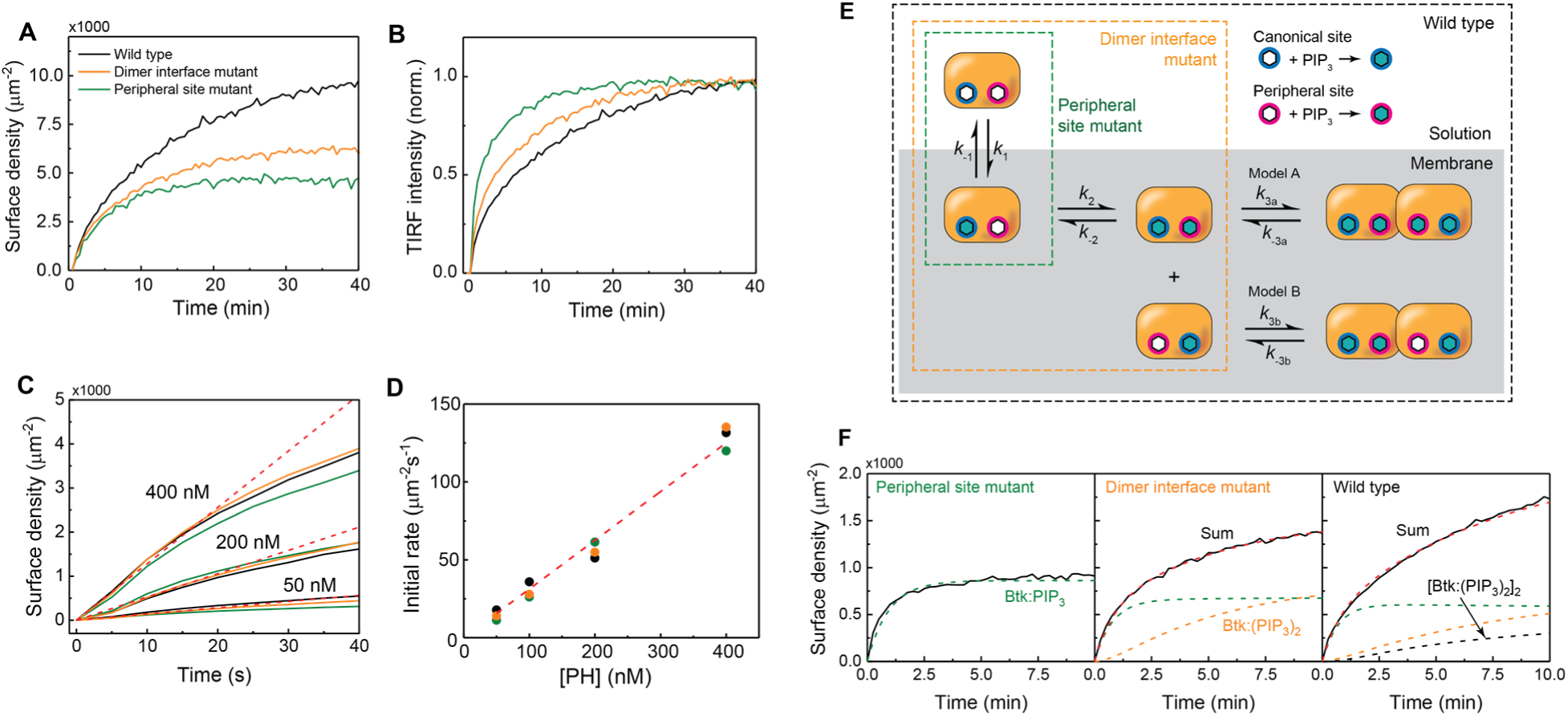
Btk PH-TH module adsorption kinetics onto PIP_3_ SLBs by TIRF microscopy. **(A)** Time-course TIRF adsorption at 40 nM measurement reveals that each PH domain construct has distinct kinetic profile. **(B)** Same data normalized to the final surface density. **(C)** The initial rates are same for all constructs. Red dotted lines show linear fits for all three constructs for a series of concentrations. **(D)** The common initial rate was estimated for all three constructs. **(E)** A simple three-step sequential kinetic model for Btk PH-TH domain is tested using mutant constructs that can undergo a subset of reactions. **(F)** The kinetic constants associated with each step were estimated by optimizing the rate laws to the TIRF adsorption measurements for different constructs.

Figure 3C shows the TIRF initial adsorption time course for each PH-TH construct at multiple concentrations. A notable feature in the kinetic adsorption profiles is that despite distinct kinetic profiles, all of the three constructs have identical initial rates, suggesting that they share a dominant initial lipid binding step. This is most likely the membrane recruitment by PIP_3_ binding at the canonical site, as the elimination of PIP_3_ binding at the canonical in the N24D/R28C mutant results in no membrane recruitment (see SI). A sequential multi-step interaction scheme was constructed, illustrated in Figure 4E. In this model, the PH-TH module is (***i***) recruited to the membrane by engaging a PIP_3_ in the canonical site, and then (***ii***) binds to another PIP_3_ in the peripheral site, followed by (***iii***) dimerization. Each PH-TH module was assumed to undergo a different subset of these processes, and this strategy allowed a sequential estimation of the rate constants, with at most two simultaneous fit parameters at a time (see SI 2). For the dimerization step, two possibilities were considered: in Model A, two PH-TH molecules were required to be anchored by both the canonical and peripheral sites to form a dimer [Btk:(PIP_3_)_2_]_2_. In Model B, on the other hand, a dually-anchored species was allowed to associate with a singly-anchored PH-TH to form a dimer [Btk:(PIP_3_)_2_]:Btk:PIP_3_. Model B includes the scenario in which a single PIP_3_ “bridges” two PH-TH modules.

In order to determine which model better represents the PH-TH module dimerization, simulated FCS data using the kinetic rate constants obtained from each model were compared to the experimental FCS data (see SI 2 for details). This comparison provides a rigorous validation for the kinetic model, as FCS is an orthogonal experiment measuring an entirely independent quantity (diffusion rate) from the TIRF measurements. Figure 5A shows the equilibrium dimer fraction as a function of the PH-TH module surface density for Models A and B, calculated using the respective kinetic rate constants. The comparison between simulated FCS based on each model with the measured data is shown in Figure 5B. It is clear that for Model B (Figure 5B, right), there is an insufficient difference between the dimer fractions for 1% and 4% PIP_3_ to account for the experimental FCS results. Model A (Figure 5B, left), however, not only captures the PIP_3_ dependence but also the overall shape of the PH-TH surface density dependence, suggesting that a PH-TH dimer is bound to at least four PIP_3_ lipids. Note that although the possibility of higher-order oligomers cannot be ruled out, dimerization following the binding of two PIP_3_ lipids to each PH-TH module is sufficient to describe these data.

**Figure 5.**
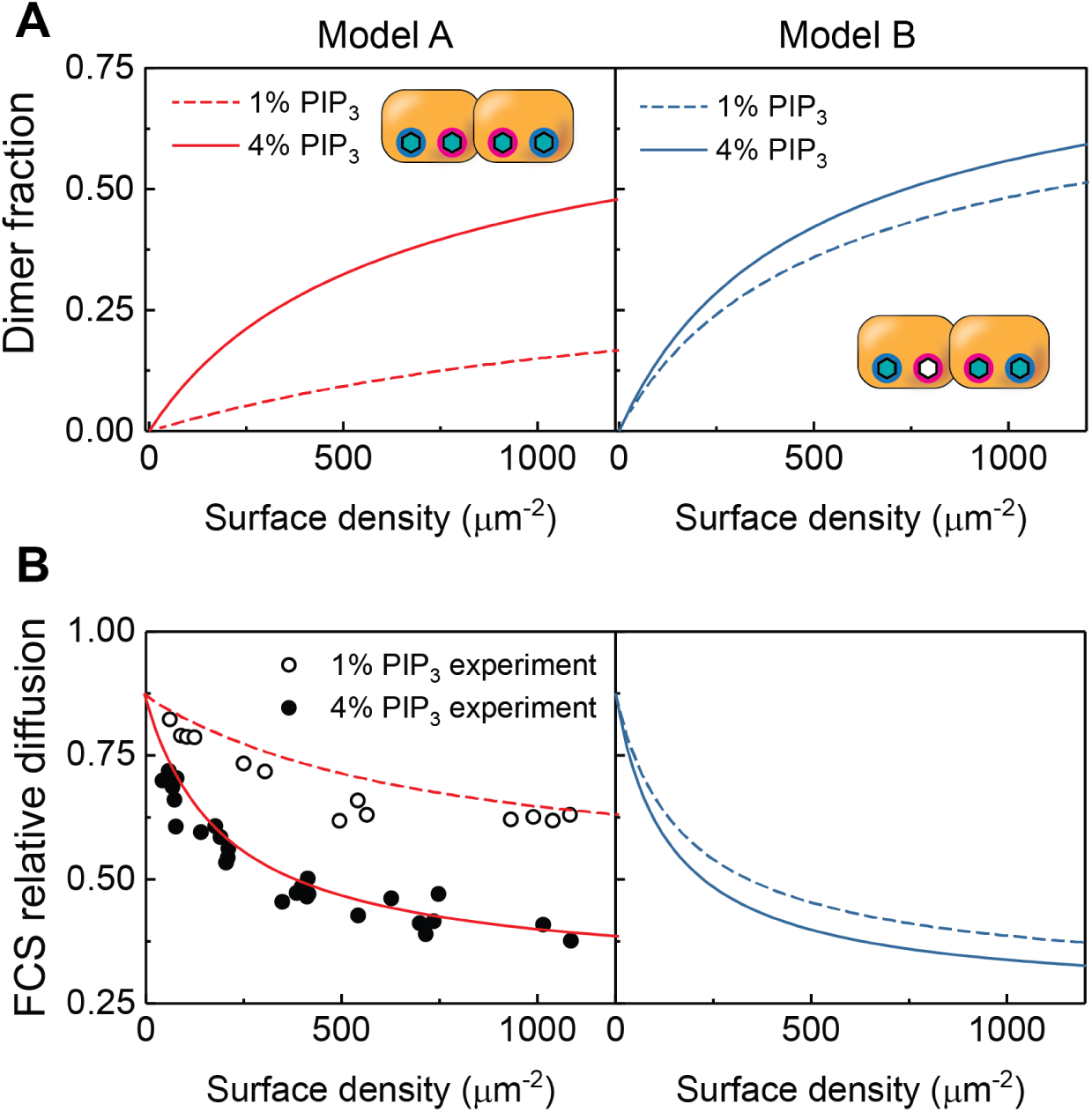
Validation of the kinetic model. **(A)** Using the kinetic rate constants obtained from the sequential lipid binding and dimerization model, the equilibrium surface density of dimers was calculated for both Model A and B cases at 1% and 4% PIP_3_. **(B)** With the results from (A), density-dependent FCS was simulated. Comparison to data suggests that in Model B, dimer case will not result in sufficient differences between 1% and 4% PIP_3_ bilayers. However, the Model A shows a good agreement with data, indicating that a Btk dimer requires four PIP_3_ binding.

These results from the adsorption kinetics modeling reveal a system adapted for switch-like PIP_3_ sensing. Because each PH-TH dimer requires four PIP_3_ molecules, the dimer concentration scales with the fourth power of the PIP_3_ surface density. This fourth-order nonlinearity creates a sharp PIP_3_ concentration threshold, which can be contrasted to a hypothetical second-order lipid sensing scenario where only one PIP_3_ binding at the canonical site is required for dimerization (Figure 6A). For the second order case (gray broken line), the dimer fraction rises gradually in response to the PIP_3_ density, with 50% dimerization observed at 1.2% PIP_3_, but with a significant dimer fraction even at a tenth of that value (10% dimer fraction at 0.12% PIP_3_). Since dimerization leads to activation, this would mean that Btk would be more vulnerable to erroneous activation due to random fluctuations in PIP_3_ concentration. Btk mitigates this problem by requiring additional PIP_3_ binding for dimerization, creating a sharper, switch-like dimerization threshold, with 50% dimerization at 3.6% PIP_3_ (black solid line). In this case, at one-tenth of that PIP_3_ concentration, the dimer fraction is only 0.4%. This ligand counting mechanism may be classified as a type of coincidence detection based on binding multiple ligand molecules.

**Figure 6.**
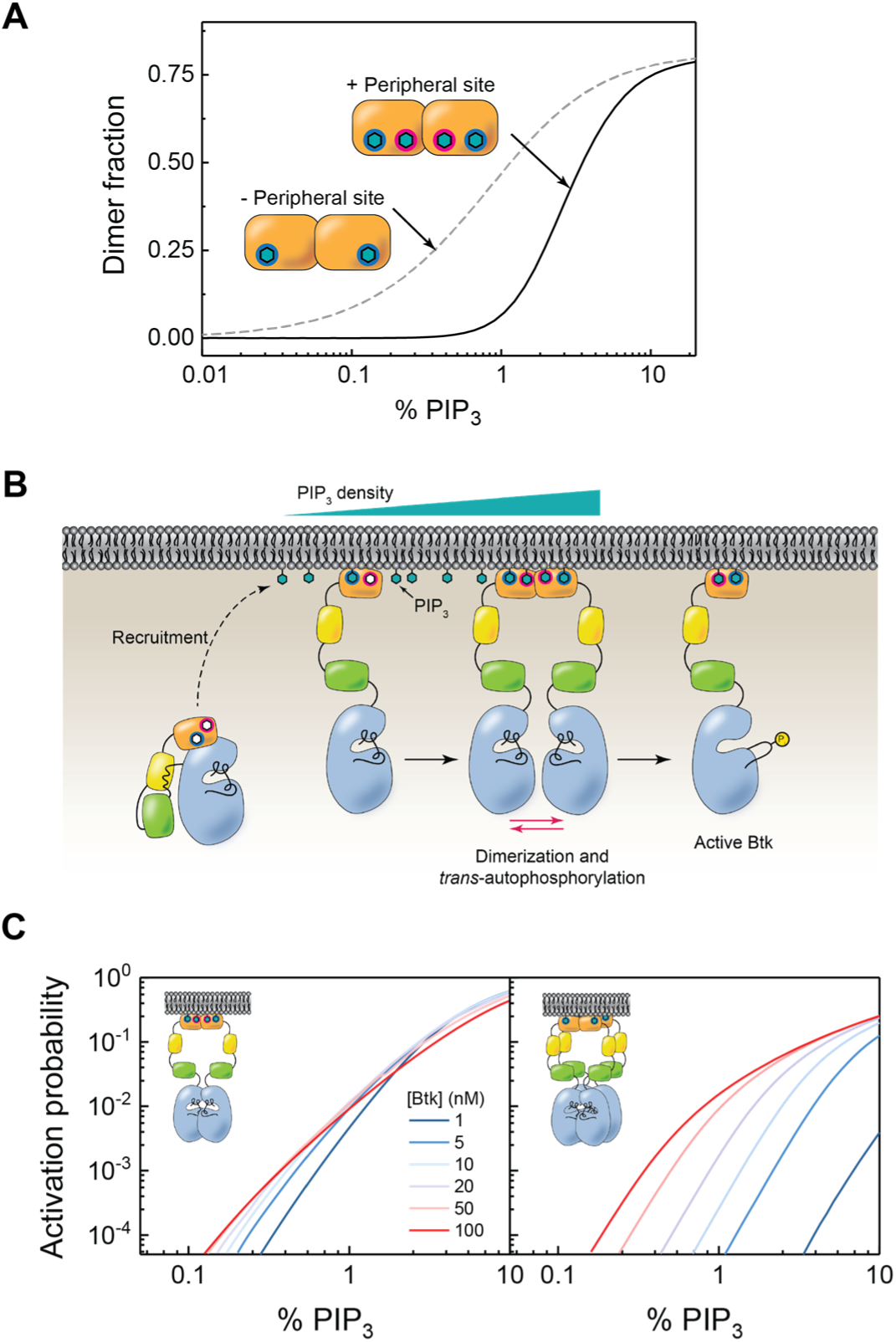
Fourth-order substrate detection by Btk. (**A**) Using the kinetic rate constants derived from the sequential kinetic model, the dimer fraction was calculated for the hypothetical case in which the PH-TH module, lacking the peripheral site, may dimerize with only one PIP_3_ (second order), and for the actual case where two are required with the peripheral site (fourth order). **(B)** In proposed activation mechanism for Btk. **(C)** An alternative method to achieve fourth-order PIP_3_ detection, single lipid binding and followed by tetramerization to activation, was considered and compared to the observed mechanism.

### PIP_3_ threshold stability to protein concentration variation

According to the model based on the FCS and adsorption data, Btk employs a system that is twofold dependent on both the ligand and protein concentrations, with an overall fourth-order detection of PIP_3_ surface density. One advantage of such a system is that it is much less sensitive to changes in protein concentration than a system that uses only 1:1 binding. To illustrate this, we considered a different scenario in which a protein forms a 1:1 complex with its ligand, in which the fourth-order ultrasensitivity to ligand concentration arises solely from tetramerization of the 1:1 complex. Figure 6C shows the activation probability of Btk (defined as the fraction of phosphorylated Btk on the membrane at equilibrium) as a function of PIP_3_ surface density for a range of Btk concentration, which was calculated based on the kinetic rate constants obtained from TIRF measurements and the catalytic rate constant of Src kinase (see SI 3 for details). In the case where tetramerization of a 1:1 PIP_3_:protein complex is required for activation, the variation of protein concentration between 1 and 100 nM significantly perturbs the threshold PIP_3_ density (Figure 6B, right). However, in the case where the PIP_3_ detection is achieved by dimerization following the binding of two PIP_3_ molecules to each protein molecule, the PIP_3_ threshold remains relatively stable over the same protein concentration range (left). This illustrates how this particular mechanism has the effect of decoupling PIP_3_ sensitivity from Btk concentration, buffering the system with respect to variations in protein expression.

### Reconstitution of Btk variants in knockout B-cells corroborate *in vitro* findings

In order to test whether these findings have a direct impact on B-cell activity in a live cell context, calcium flux in a Btk-knockout chicken B-cell line, DT40, was measured with various mutations present. Indo-1 calcium indicator measurements these mutant Btk, normalized for expression, are shown in Figure 7. Wild-type Btk shows expression level-dependent calcium flux, demonstrating that the B-cell activation is dependent on the Btk concentration (Figure S4). The negative control, the kinase dead mutant D521N, shows a clear reduction of calcium flux, while the positive control, the gain-of-function mutant E41K (whose PH-TH module has been modified to bind PIP_2_) shows saturating activation, even at a lower Btk concentration than the wild-type (Dingjan et al., 1998). For both the dimer interface and peripheral site mutants, there is a comparable decrease of calcium flux to the kinase dead mutant. This suggests that Btk membrane recruitment through both the canonical and peripheral sites, as well as its dimerization, contributes to the overall activation probability of Btk. In the absence of dimerization, Btk may be activated by other kinases or through slow *trans*-autophosphorylation. However, the membrane dwell time would be shortened, and Btk would be less likely to encounter PLCγ2. Therefore, robust B-cell signaling is dependent on the ability of the PH-TH module to bind PIP_3_ in both binding sites as well as undergo dimerization.

**Figure 7.**
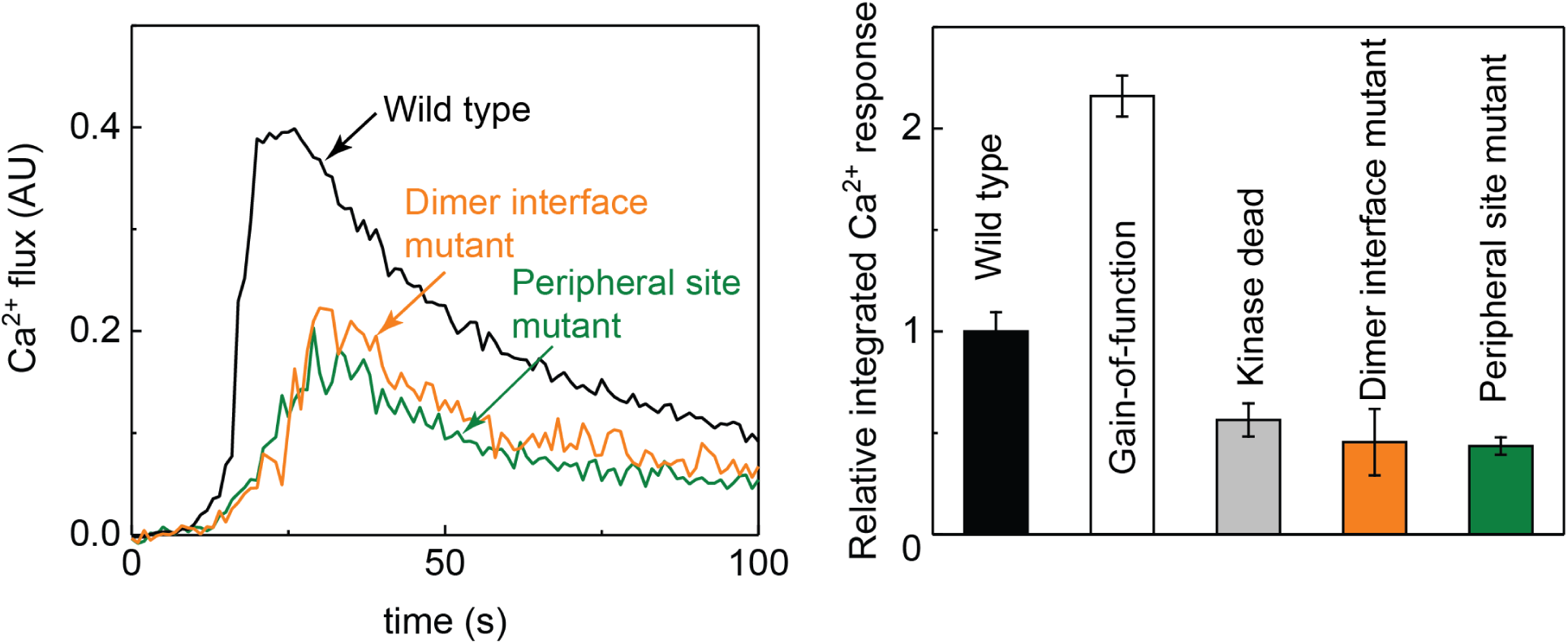
Calcium response by Btk variants in B-cells. **(A)** Btk-deficient chicken B-cells were transfected with variants of Btk and activated with BCR antibody added at 0 s, and monitored for calcium flux. **(B)** Relative integrated calcium response for WT, gain-of-function (E41K), kinase dead (D521N), dimer interface mutant, and peripheral site mutant.

## CONCLUSIONS

Activation by dimerization-facilitated *trans*-autophosphorylation is a common feature of protein kinases. Indeed, the adventitious dimerization of tyrosine kinases leads to improper activation and cancer, as exemplified by the BCR-Abl fusion protein (Hantschel et al., 2003). The ability of non-receptor tyrosines kinases such as Src and Abl to activate through dimerization and *trans*-autophosphorylation is well established, but the natural mechanisms that promote dimerization are still not well understood (Lamontanara et al., 2014; Xu et al., 2015). In this regard, the unexpected discovery that Btk can be activated in solution by the binding of IP_6_ to the PH-TH module in a way that is consistent with the crystallographically observed Saraste dimer was intriguing, but there was no direct evidence that dimerization was in fact responsible for the observed activation (Wang et al., 2015). In this work, we have now established that dimerization of the Btk PH module can be observed on reconstituted membrane surfaces. By measuring diffusion and membrane binding kinetics, we show that PIP_3_ binding at peripheral site of the PH-TH module is necessary for the dimerization, and this requirement gives rise to a sharp threshold for activation.

A recent study using NMR and hydrogen-deuterium exchange mass spectrometry (HDX-MS) has shown Btk primarily exists in the autoinhibited form in solution, and in order for the PH-TH module to engage PIP_3_, this conformation must be released, in agreement with earlier SAXS experiments (Joseph et al., 2017; Marquez et al., 2003). According to this study, the β3-β4 loop of the PH-TH module, which includes the dimerization and peripheral binding sites, is involved in allosterically regulating the autoinhibited form of Btk by blocking the activation loop of the kinase domain. This is consistent with previous work which showed increased basal activity of Btk in the absence of the PH-TH module (Wang et al., 2015). A crystal structure of a construct of Btk containing the PH-TH module and the kinase domain, with the SH2 and SH3 domains deleted, showed a different conformation, with the PH-TH module docked on top of the N-terminal lobe of the kinase domain, away from the active site (Wang et al., 2015). In this structure, the active site is unimpeded, and the PH-TH module forms a Saraste dimer in the crystal. This suggests that one role for dimerization of the PH-TH module might be to release blockage of the kinase active site, allowing phosphorylation of the activation loop of the kinase domain by *trans*-autophosphorylation or by another kinase. Overall, the PH-TH module displays multiple functions including recruitment to the membrane, detection of the PIP_3_ level, and cooperative PIP_3_ binding leading to dimerization.

A key consequence of the additional PIP_3_ binding at the peripheral site is that it makes Btk extremely sensitive to the PIP_3_ density, a tightly regulated but dynamic signal that influences numerous signaling pathways. Activation by *trans*-autophosphorylation is a positive feedback mechanism for a switch-like response to signal (Shinohara et al., 2014). By requiring the peripheral site to bind a PIP_3_ for activation by dimerization, the Btk PH-TH module creates an even sharper PIP_3_ concentration threshold for activation. The mutational studies in B-cells also supported the prediction that these protein-membrane interactions are critical in B-cell activation. Together, they provide detailed insights to how B-cells may be instituting a minimalist coincidence detection, and more broadly, how lipid-binding domains have evolved to enhance the integrity of cellular signal transduction. Considering the diversity and omnipresence of lipid-binding domains in the proteome, it is likely that this simple and effective mechanism is not unique to Btk and present in other signaling proteins as well.

Another question that emerges from this work is whether any of the newly discovered properties of Btk PH-TH module can be found in PH domains in other proteins (Karandur et al., 2017; Yamamoto et al., 2016). A particularly interesting case is Itk, another Tec family kinase that has Btk’s role in T-cell signaling. Sequence alignment indicates that Itk lacks key residues in the peripheral site and the dimer interface, alluding its operation would not be similar to Btk’s. These differences may provide insights to how T-cells and B-cells have adapted to different roles in the course of the development of the immune system.

## METHODS

### Protein purification

The purification protocol for the PH-TH module constructs used in this study have been published previously (Wang et al., 2015). Briefly, pET-28 plasmid containing residues 1-172 of bovine Btk fused to C-terminal eGFP or mCherry and N-terminal His6 was transformed into BL21(DE3) cells with GroES/GroEL and YopH. After growth and lysis, the proteins were purified by a Ni-NTA affinity chromatography, and the His tag was removed, followed by size exclusion chromatography.

### Supported lipid bilayers (SLB)

Planar SLBs of varying compositions consisting primarily of DOPC (Avanti Polar Lipids) were prepared for fluorescence microscopy. The detailed protocol has been published previously (Lin et al., 2010). Briefly, SLBs were formed by rupturing small unilamellar vesicles, prepared by probe sonication, on glass substrates cleaned by piranha etch. Other bilayer components such as PI(3,2,4)P_3_ (PIP_3_), PI(4,5)P_2_ (PIP_2_), DOPS were also purchased from Avanti. A trace amount (0.005 mol %) of Texas Red DHPE (ThermoFisher Scientific) was also included. The bilayers were prepared in ibidi sticky-slide VI 0.4 microfluidic chambers (ibidi GmbH). The final buffer (pH 7.4) was composed of the following: 40 mM HEPES, 100 mM NaCl, 10 mM BME, 0.1 mg/mL casein.

### Fluorescence correlation spectroscopy (FCS) and FRET

Dual-color FCS measurements were performed on a home-built confocal system integrated into an inverted microscope. The experimental methods have been published previously (Chung et al., 2016). The light source was a pulsed (100 ps) supercontinuum laser (NKT Photonics). The average excitation power for a typical FCS measurement was 0.5 and 1 μW for 488 and 568 nm, respectively. The signals were collected by the objective and passed a 50-μm pinhole, then detected by avalanche photodiode detectors (Hamamatsu), and processed by a hardware correlator (Correlator.com). The focus radii, calibrated with bilayers containing known amounts of fluorescent lipids, were 0.20 ± 0.01 μm and 0.22 ± 0.01 μm for 488 and 568 nm, respectively. For FRET, fluorescence lifetime of the donor fluorophore (eGFP) in the presence of an acceptor fluorophore (mCherry) by TCSPC. The measurements were performed on the same system but the signals were processed by a TCSPC card (PicoQuant). The resulting photon arrival time histograms were fit to a single exponential decay with lifetime *τ*.

### Adsorption kinetics

The adsorption and desorption kinetics of eGFP-labeled Btk PH-TH domain constructs were obtained by bulk total internal reflection fluorescence (TIRF) microscopy (Nikon Ti2000), with objective-type TIRF illumination (NA 1.49). The excitation source was a 488 nm diode laser (Coherent Cube), and the camera was Andor iXon EMCCD (Andor Technology). The analysis of kinetic time courses simulated by numerically solving coupled kinetic equations in MATLAB. See SI for details.

### Btk knockout B-cell calcium activity assay

Btk-deficient DT40 cells were transiently transfected with 15 μg DNA of pEGFP-N1-nBtk (wild type or mutant) by electroporation (Biorad). The cells were then washed in cell loading media (CLM) containing 1 mM Ca^2+^, 1 mM Mg^2+^, 1% FBS and loaded with 1mg/mL indo-1 AM (Life Technologies) and 1mM probenecid for 30 minutes at 37°C. Then, they were washed twice with CLM, resuspended in 1 mL CLM, and rested for 15 minutes prior to analysis. Calcium flux was monitored by flow cytometry (BD LSRFortessa) before and after stimulation with 1:1000 dilution of anti-chicken BCR monoclonal antibody M4 or 1 μM Ionomycin.

## ACKNOWLEDGMENTS

We would like to acknowledge Angelica Gonzalez-Sanchez and Han Li for pilot experiments. We would also like to thank Dr. Neel Shah, Dr. Scott D. Hansen, and Kiera Wilhelm for critical reading of the manuscript. The primary support for this work was provided by the National Health Institute (NIH), under P01AI091580. Additional support was provided by the National Cancer Institute, NIH, under U01CA202241.

## AUTHOR CONTRIBUTIONS

Conceptualization, J.K.C., L.M.N., Q.W., J.K., J.T.G.; Methodology, J.K.C., J.T.G.; Investigation and Validation, J.K.C., L.M.N., T.A.K.; Writing – Original Draft, J.K.C., L.M.N., J.K., J.T.G.; Writing – Review & Editing, all; Supervision and Funding Acquisition, A.W., J.K., J.T.G

