## Supplementary Materials for "Switch-like activation of Bruton’s tyrosine kinase by membrane-mediated dimerization"

### 1. TIRF calibration

For the kinetic adsorption measurements, the surface density was measured by bulk TIRF, which was calibrated against FCS density measurements. The dimer interface mutant was used for the calibration to avoid complications arising from dimerization. For the experimental range of 0 – 1000 molecules per $\mu m^2$, the TIRF intensity was linear with the surface density.

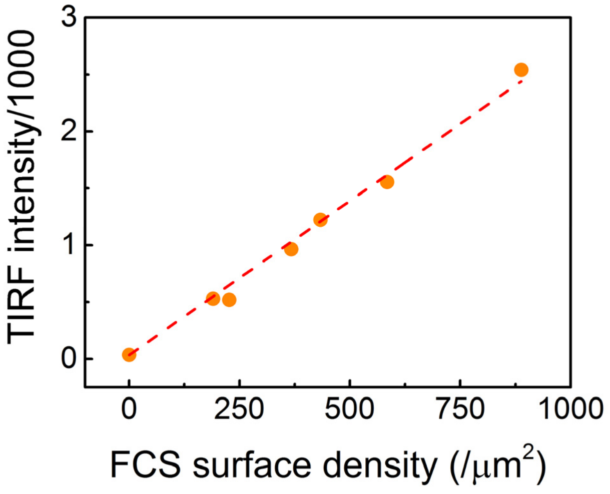

**Figure S1. TIRF calibration** TIRF intensity was calibrated against surface density measured by FCS with the dimer interface mutant.

### 2. The canonical site mutant

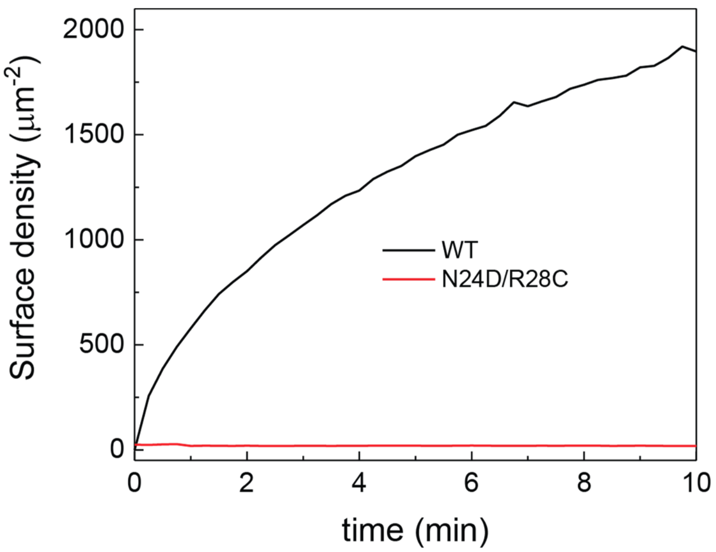

**Figure S2. Membrane adsorption of the canonical site mutant, N24D/R28C.** When the $PIP_3$ binding at the canonical site is disrupted by N24D/R28C mutation, there is negligible membrane recruitment, suggesting that the first kinetic step seen in the wild type is due to the specific $PIP_3$ binding at the canonical site. [$PIP_3$] = 4%, [WT] = 10 nM, [N24D/R28C] = 20 nM.

### 3. Estimating kinetic rate constants from TIRF adsorption

We have three Btk PH domain constructs: the wild type, dimer interface mutant, and the peripheral site mutant. Each construct has a distinct adsorption kinetic profile (Figure 4A). We will use this adsorption kinetics data to construct a model for Btk PH domain binding to the $PIP_3$ membrane followed by dimerization.

The simplest model for the wild type, in three steps, is:

1. $PIP_3$ binding at the canonical site
2. $PIP_3$ binding at the peripheral site
3. Dimerization

First two steps are simply:

(1) Btk + $PIP_3 \leftrightarrow$ Btk:$PIP_3$ ($k_1/k_{-1}$ fwd/rev rate constants)

(2) Btk:$PIP_3$ + $PIP_3 \leftrightarrow$ Btk:$(PIP_3)_2$ ($k_2/k_{-2}$ fwd/rev rate constants)

Two possibilities for the third step, dimerization, were considered. In the first case, each PH domain will be required to bind to two $PIP_3$ lipids in order to dimerize.

(3a) 2Btk:$(PIP_3)_2 \leftrightarrow Btk_2:(PIP_3)_4$ ($k_{3a}/k_{-3a}$ fwd/rev rate constants)

In the other case, only one PH domain is required to be bound to two $PIP_3$:

(3b) Btk:$(PIP_3)_2$ + Btk:$PIP_3 \leftrightarrow Btk_2:(PIP_3)_3$ ($k_{3b}/k_{-3b}$ fwd/rev rate constants)

**Model A** will follow the steps (1) → (2) → (3a), and **Model B** will follow (1) → (2) → (3b).

We will use the simplified notation

$B$ = Btk

$P$ = $PIP_3$

Then, the mass action rate laws are:

**Model A**:

$$d[B]/dt = -k_1[B][P] + k_{-1}[BP]$$
$$d[P]/dt = -k_1[B][P] + k_{-1}[BP] - k_2[BP] + k_{-2}[BP_2]$$
$$d[BP]/dt = k_1[B][P] - k_{-1}[BP] - k_2[P][BP] + k_{-2}[BP_2]$$
$$d[BP_2]/dt = k_2[BP][P] - k_{-2}BP_2 - 2k_{3a}[BP_2]^2 + 2k_{-3a}[B_2P_4]$$
$$d[B_2P_4]/dt = k_{3a}[BP_2]^2 - k_{-3a}[B_2P_4]$$

**Model B:**

$$d[B]/dt = -k_1[B][P] + k_{-1}[BP]$$
$$d[P]/dt = -k_1[B][P] + k_{-1}[BP] - k_2[BP] + k_{-2}[BP_2]$$
$$d[BP]/dt = k_1[B][P] - k_{-1}[BP] - k_2[P][BP] + k_{-2}[BP_2] - k_{3b}[BP][BP_2] + k_{-3b}[B_2P_3]$$
$$d[BP_2]/dt = k_2[BP][P] - k_{-2}BP_2 - k_{3b}[BP][BP_2] + k_{-3b}[B_2P_3]$$
$$d[B_2P_3]/dt = k_{3b}[BP][BP_2] - k_{-3b}[B_2P_3]$$

The kinetic constants will be estimated by fitting the TIRF data with these two models. This was accomplished by sequentially fitting each mutant constructs, as illustrated in Figure 4E.

For the peripheral site mutant the reaction cannot proceed past the first step, meaning $k_2 = k_3 = 0$. The coupled kinetic equations are then reduced to

$$d[B]/dt = -k_1[B][P] + k_{-1}[BP]$$
$$d[P]/dt = -k_1[B][P] + k_{-1}[BP]$$
$$d[BP]/dt = k_1[B][P] - k_{-1}[BP]$$

Here, there are only two kinetic constants, $k_1$ and $k_{-1}$. $k_1$ can be independently estimated from the initial velocity. $k_1$ is same for all constructs, which is consistent with our assumptions about the sequential steps and independent binding.

The system involves both 2-dimensional (membrane-bound) and 3-dimensional (solution phase) species, which may complicate units. For simplicity, absolute number of molecules (in pmol, $6.022 \times 10^{-11}$) instead of concentration was used to represent the amount of each species in the experimentally defined conditions. With this choice of units, the forward reaction rate constants are in $\text{pmol}^{-1}\text{s}^{-1}$ and reverse rate constants are in $\text{s}^{-1}$. The surface density of membrane-bound species measured by FCS-calibrated TIRF was likewise converted to pmol. Fitting the peripheral site mutant adsorption data with $k_1$ fixed at the measured value, $k_{-1}$ was estimated by least square optimization.

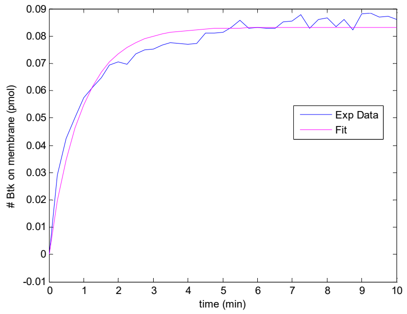

For the dimer interface mutant, $k_3 = 0$ as it cannot dimerize. The rate equations are:

$$d[B]/dt = -k_1[B][P] + k_{-1}[BP]$$
$$d[P]/dt = -k_1[B][P] + k_{-1}[BP] - k_2[BP] + k_{-2}[BP_2]$$
$$d[BP]/dt = k_1[B][P] - k_{-1}[BP] - k_2[P][BP] + k_{-2}[BP_2]$$
$$d[BP_2]/dt = k_2[BP][P] - k_{-2}BP_2$$

Here, there are four kinetic constants, $k_1$, $k_{-1}$, $k_2$, and $k_{-2}$. $k_1$ and $k_{-1}$ are fixed at the values obtained from the peripheral site mutant data in the previous step, so only $k_2$ and $k_{-2}$ need to be estimated. In this case, the sum $[BP] + [BP_2]$ was optimized to the dimer interface mutant adsorption data, since both species contribute to the TIRF intensity equally.

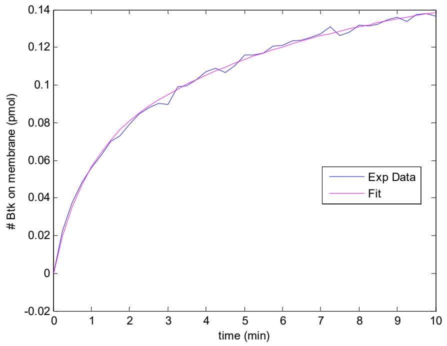

For the last step, the same approach was used to estimate $k_{3a}$, $k_{-3a}$ with Model A, and $k_{3b}$, $k_{-3b}$ with Model B. Here, the membrane-bound species present in the system are BP, $BP_2$, and either of the dimeric species $B_2P_3$ or $B_2P_4$. Since the dimeric species are expected to contribute about twice as much to the TIRF intensity, the sum $[BP] + [BP_2] + 2[B_2P_3]$ and $[BP] + [BP_2] + 2[B_2P_4]$, for Model A and B, respectively, were optimized to the wild type adsorption data.

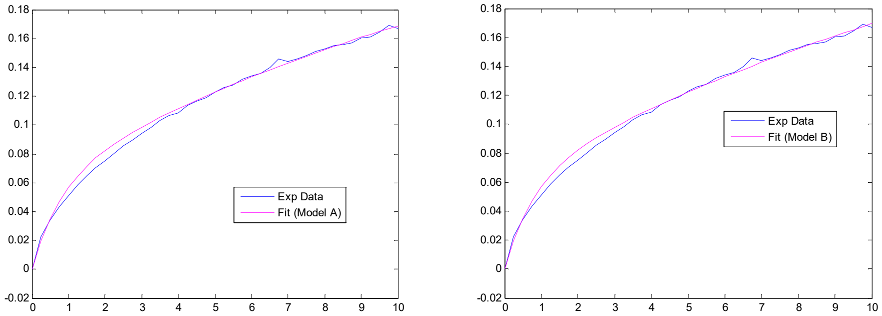

The following table summarizes the estimated kinetic constants:

| Reaction | Forward rate constant ($pmol^{-1}s^{-1}$) | Reverse rate constant ($s^{-1}$) |
|---|---|---|
| $B + P \leftrightarrow BP$ | $k_1 = 4.7 \times 10^{-4}$ | $k_{-1} = 1.5 \times 10^{-3}$ |
| $BP + P \leftrightarrow BP_2$ | $k_2 = 7.8 \times 10^{-4}$ | $k_{-2} = 3.1 \times 10^{-3}$ |
| $2BP_2 \leftrightarrow B_2P_4$ | $k_{3a} = 1.0$ | $k_{-3a} = 5.0 \times 10^{-2}$ |
| $BP_2 + BP \leftrightarrow B_2P_3$ | $k_{3b} = 2.5 \times 10^{-2}$ | $k_{-3b} = 1.5 \times 10^{-4}$ |

In more familiar units,

| Reaction | Forward rate constant | Reverse rate constant ($s^{-1}$) |
|---|---|---|
| $B + P \leftrightarrow BP$ | $2.8 \times 10^{-3}\ nM^{-1}s^{-1}$ | $1.5 \times 10^{-3}$ |
| $BP + P \leftrightarrow BP_2$ | $7.4 \times 10^{6}\ \mu m^2\ s^{-1}$ | $3.1 \times 10^{-3}$ |
| $2BP_2 \leftrightarrow B_2P_4$ | $9.4 \times 10^{-5}\ \mu m^2\ s^{-1}$ | $5.0 \times 10^{-2}$ |
| $BP_2 + BP \leftrightarrow B_2P_3$ | $2.4 \times 10^{-6}\ \mu m^2\ s^{-1}$ | $1.5 \times 10^{-4}$ |

### 4. Calculating the activation probability for dimerization vs. tetramerization

This section will detail activation probability shown in Figure 6. For the dimerization case, the model is

(1) $\mathrm{Btk + PIP_3 \underset{k_{-1}}{\overset{k_1}{\rightleftharpoons}} Btk{:}PIP_3}$

(2) $\mathrm{Btk{:}PIP_3 + PIP_3 \underset{k_{-2}}{\overset{k_2}{\rightleftharpoons}} Btk{:}(PIP_3)_2}$

(3) $\mathrm{2Btk{:}(PIP_3)_2 \underset{k_{-3}}{\overset{k_3}{\rightleftharpoons}} Btk_2{:}(PIP_3)_4 \xrightarrow{k_{cat}} 2\mathit{p}Btk{:}(PIP_3)_2}$

(4) $\mathrm{\mathit{p}Btk{:}(PIP_3)_2 \underset{k_2}{\overset{k_{-2}}{\rightleftharpoons}} \mathit{p}Btk{:}PIP_3 + PIP_3}$

(5) $\mathrm{\mathit{p}Btk{:}PIP_3 \underset{k_1}{\overset{k_{-1}}{\rightleftharpoons}} \mathit{p}Btk + PIP_3}$

And for the tetramerization case,

(1) $\mathrm{Btk + PIP_3 \underset{k_{-1}}{\overset{k_1}{\rightleftharpoons}} Btk{:}PIP_3}$

(2) $\mathrm{4Btk{:}PIP_3 \underset{k_{-3}}{\overset{k_3}{\rightleftharpoons}} (Btk{:}PIP_3)_4 \xrightarrow{k_{cat}} 4\mathit{p}Btk{:}PIP_3}$

(3) $\mathrm{\mathit{p}Btk{:}PIP_3 \underset{k_1}{\overset{k_{-1}}{\rightleftharpoons}} \mathit{p}Btk + PIP_3}$

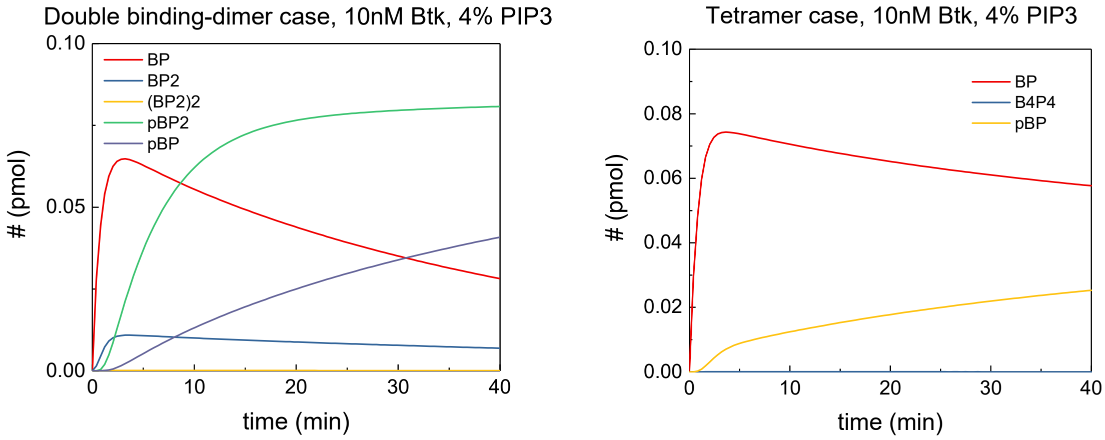

The time course for different species for the double binding-dimer case, and the tetramer case for 40 minutes are shown in the above figures. The kinetic rate constants calculated from Model A and $k_{cat}$ from Src (Wang et al., 2015) were used. Then, the activation probability was defined as the number of membrane-bound phosphorylated Btk divided by all membrane-bound Btk at equilibrium. In other words, for the dimerization case,

$$P_d = ( [pBtk{:}PIP_3] + [pBtk{:}(PIP_3)_2])/ ( [pBtk{:}PIP_3] + [pBtk{:}(PIP_3)_2] + [Btk{:}PIP_3] + [Btk{:}(PIP_3)_2])$$

And for the tetramerization case,

$$P_t = [pBtk{:}PIP_3]/ ([pBtk{:}PIP_3] + [Btk{:}PIP_3])$$

Same calculations were performed for a range of $PIP_3$ surface densities to obtain Figure 6C.

### 5. Effects of other anionic lipids

To confirm that the density dependent dimerization effect observed on $PIP_3$ is $PIP_3$ specific, alternative anionic lipids were used to perform FCS on supported lipid bilayers. In the case of PS, the lipid shows no enhancement of dimerization in the 1% $PIP_3$ case. When combining $PIP_2$ with $PIP_3$, an intermediate level of dimerization can be achieved, however the presence of $PIP_2$ cannot recapitulate the dimer populations observed for 4% $PIP_3$, even at very high Btk PH-TH surface density. It should be noted that Btk PH-TH does not associate to bilayers containing only $PIP_2$, in the absence of $PIP_3$.

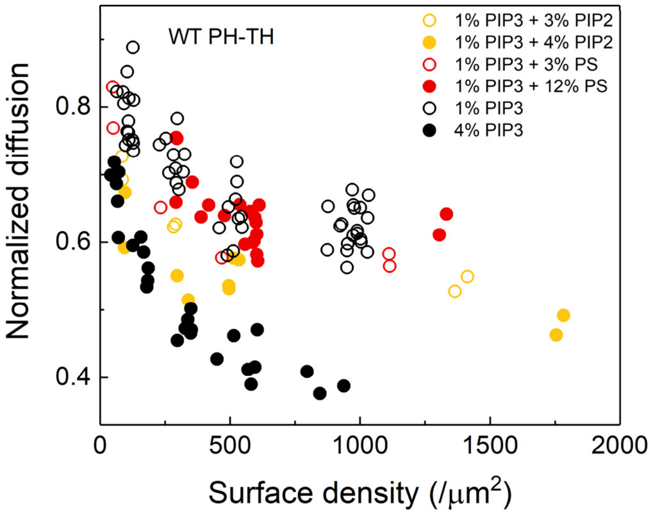

**Figure S3. Effects of other anionic lipids.** Btk PH-TH module dimerization for various lipid bilayer compositions, including $PIP_3$.

### 6. Effect of $IP_6$

As it was not previously known whether the peripheral site is specific for $IP_6$ alone, experiments were performed to confirm that $IP_6$ does not enhance dimerization of Btk PH-TH on $PIP_3$ containing bilayers. This is true for the peripheral site mutant as well. In fact, $IP_6$ seems to directly compete with $PIP_3$ and will compete Btk PH-TH off the membrane at high enough concentrations.

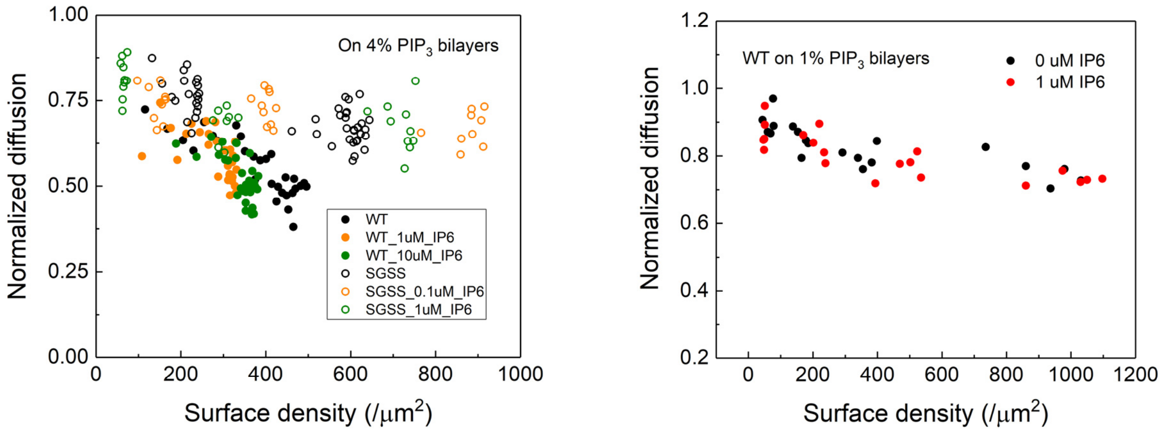

**Figure S4. Effect of IP6 on PH-TH module dimerization.** (*Left*) On 4% $PIP_3$ bilayers, the wild type PH-TH and the peripheral site mutant shows identical dimerization behavior with or without $IP_6$. (*Right*) On 1% PIP3 bilayers, the presence of $IP_6$ does not promote dimerization.

### 7. Expression level-dependent B-cell activation with Btk variants

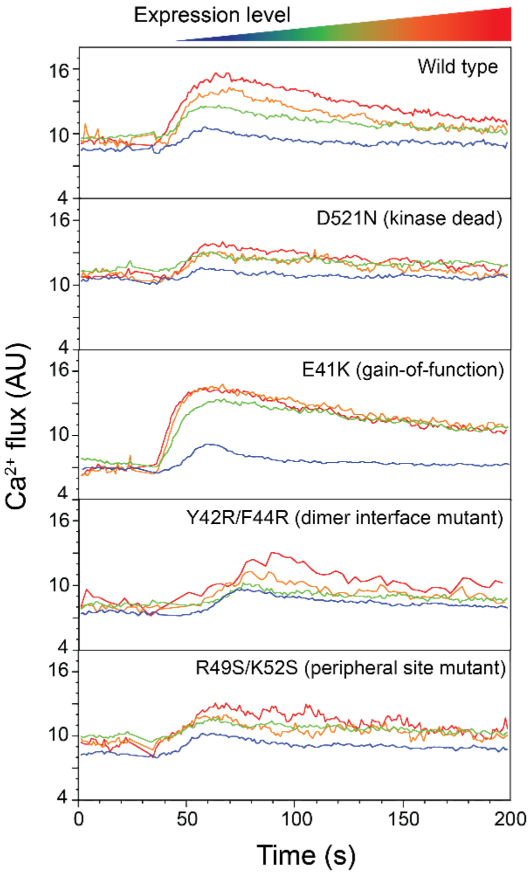

**Figure S5. Expression level-dependent B-cell activation for Btk variants.** The calcium flux upon activation by anti-BCR antibody are shown for wild type, D521N (kinase dead, negative control), E41K (gain-of-function, positive control), Y42R/F44R (dimer interface mutant), and R49S/K52S (peripheral site mutant). The dimer interface and peripheral site mutants show calcium flux comparable to that of D521N, demonstrating that both dimerization and the peripheral $PIP_3$ binding are important for Btk activation.
